# Massively Parallel Microwire Arrays Integrated with CMOS chips for Neural Recording

**DOI:** 10.1101/573295

**Authors:** Abdulmalik Obaid, Mina-Elraheb Hanna, Yu-Wei Wu, Mihaly Kollo, Romeo Racz, Matthew R Angle, Jan Müller, Nora Brackbill, William Wray, Felix Franke, E.J. Chichilnisky, Andreas Hierlemann, Jun B Ding, Andreas T Schaefer, Nicholas A Melosh

**Affiliations:** Department of Materials Science and Engineering, Stanford University, Stanford, CA, USA; Department of Neurosurgery, Stanford University School of Medicine, Stanford, CA, USA; Neurophysiology of Behaviour Laboratory, Francis Crick Institute, London, UK; Department of Neuroscience, Physiology & Pharmacology, University College London, London, UK; Paradromics Inc., Austin, TX, USA; Department of Biosystems Science and Engineering, ETH Zurich, Basel, Switzerland; Department of Physics, Stanford University, Stanford, California, USA; Departments of Neurosurgery and Ophthalmology, Stanford University, Stanford, California, USA

## Abstract

Multi-channel electrical recordings of neural activity in the brain is an increasingly powerful method revealing new aspects of neural communication, computation, and prosthetics. However, while planar silicon-based CMOS devices in conventional electronics scale rapidly, neural interface devices have not kept pace. Here, we present a new strategy to interface silicon-based chips with three-dimensional microwire arrays, providing the link between rapidly-developing electronics and high density neural interfaces. The system consists of a bundle of microwires mated to large-scale microelectrode arrays, such as camera chips. This system has excellent recording performance, demonstrated via single unit and local-field potential recordings in isolated retina and in the motor cortex or striatum of awake moving mice. The modular design enables a variety of microwire types and sizes to be integrated with different types of pixel arrays, connecting the rapid progress of commercial multiplexing, digitisation and data acquisition hardware together with a three-dimensional neural interface.

## Introduction

Neural activity occurs within interconnected populations of neurons operating over a range of length scales and spatial locations throughout the brain. Recording sufficient numbers of neurons at natural timescales and spatial distributions is one of the foremost challenges for improved understanding of how neuronal ensembles operate in different behavioural states^1–3^. Optical methods are increasingly used in studies of network dynamics *in vivo*, as they permit the monitoring of activity over a large area from the same layer of brain tissue^4–7^. While improving, current optical techniques are limited in sampling rate, obscuring fine temporal patterns which have been found to carry significant information in neuronal population codes^8–10^. Furthermore, they are inherently limited to recording from superficial structures or require the use of e.g. microendoscope probes or surgical resection of overlying tissue to provide access to deeper brain regions^11–14^.

A number of innovations have been made for electrical recording using flexible materials on the surface of the brain, and planar probes have seen rapid development based on silicon processing techniques^15–17^. However, recording from volumetrically distributed sites at scale has remained challenging due to the inherently two-dimensional nature of these devices. Since many brain areas (e.g., neocortex, hippocampus, olfactory bulb) are organized in strata, sampling broad horizontal layers over large areas would be highly beneficial, as demonstrated by imaging experiments^18,19^.

Microelectrodes and Si microarrays (Utah Arrays^20^) have long been the standard for high speed, distributed recording electrodes, yet are limited in channel count due to connectorization, volumetric displacement and tissue damage. At the same time, CMOS-based electronics continues to evolve at a rapid pace for slice and culture recordings^21–26^, yet few of these technological improvements make their way to *in vivo* neuroscience^27^. Apart from individual electrodes, scaling has been frustrated by the engineering challenge for connectors, amplification, and digitization with integrated active electronics^16,28^, posing noise and temperature challenges^14^.

Here, we report a new strategy to take advantage of the scalability and electronic processing power of CMOS-based devices combined with a three-dimensional neural interface. The core concept is shown in Fig 1a, consisting of a bundle of insulated microwires perpendicularly mated to a large-scale CMOS amplifier array, such as a pixel array found in commercial camera or display chips. While microwires have low insertion damage and excellent electrical recording performance^30–32^, they have been difficult to scale because they require individual mounting and connectorization^33,34^. By arranging them into bundles, we control the spatial arrangement and three-dimensional structure of the distal (neuronal) end, with a robust parallel contact plane on the proximal side mated to a planar pixel array.

**Figure 1:**
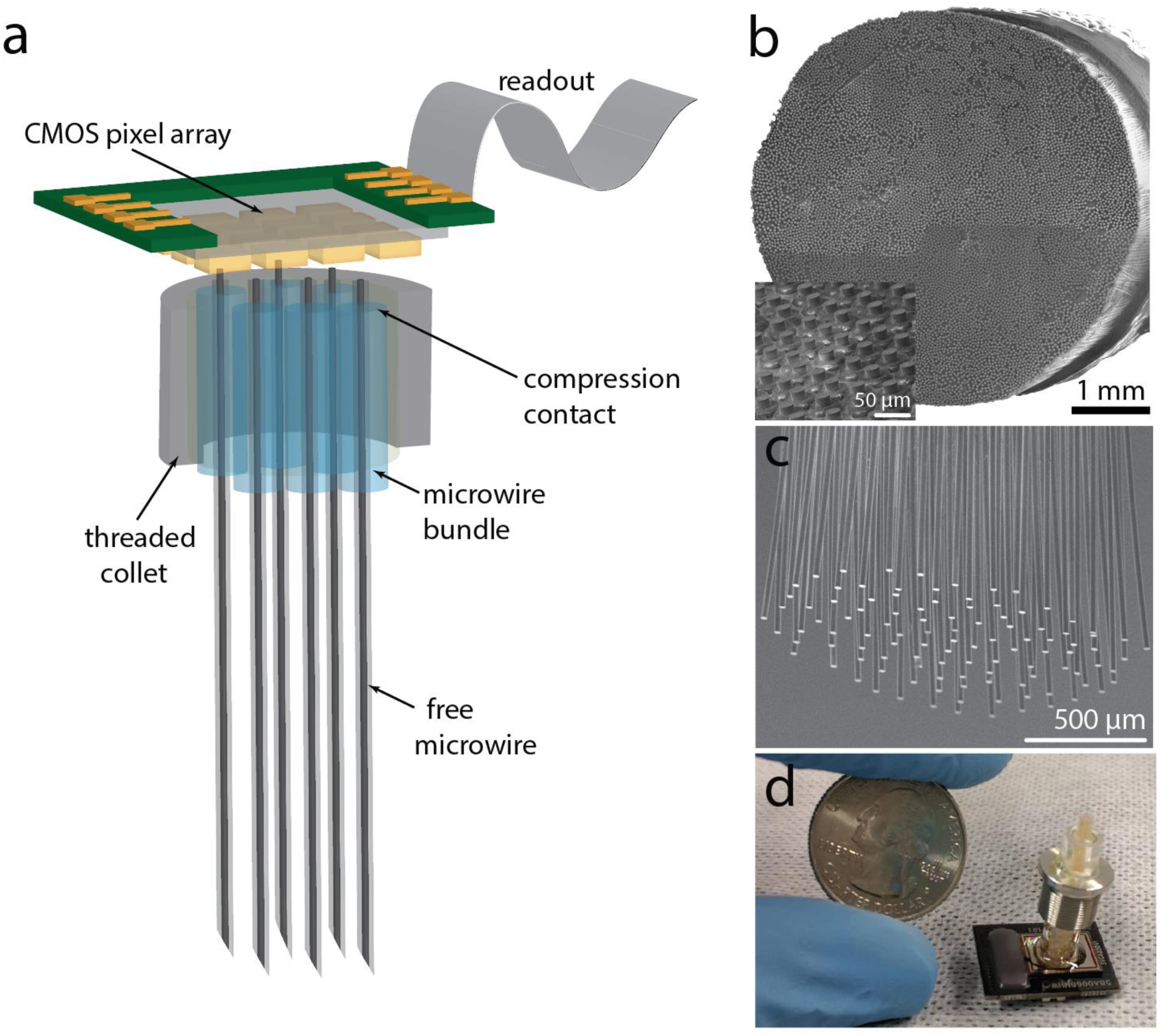
Neural bundle design. **a**, Schematic of the CMOS chip integrated with the microwire bundle. The bundle consists of a proximal (chip) end, **b**, designed for contact to the CMOS pixels, and a distal (brain) end, **c**, designed to record in tissue. The proximal end has partially exposed metal wires to contact the chip, while the distal end wires are separated to limit tissue damage upon insertion. **d**, A bundle of 800 microwires spaced 100 µm apart, with a device form factor less than 0.6 cm wide appropriate for small animal studies.

This architecture provides an array of microwires over the brain surface, akin to a Utah array^20^, rather than along a single recording plane, such as with Si or polymer shank probes^16,35^. Moreover, the insertion depth of the microelectrodes can be shaped to accommodate specific spatial distributions. The modular nature of the design allows a wide array of microwire types and size to be mated to different CMOS chips. The density of the microwires for the proximal (chip) end (Fig 1b) and the distal (brain) end (Fig 1c) can be modulated independently (Fig S3h), allowing the wire-to-wire spacing to be tailored as desired. We thus link the rapid progress and power of commercial CMOS multiplexing, digitization and data acquisition hardware together with a bio-compatible, flexible and sensitive neural interface array.

## Results and Discussion

### Microwire Bundles

The microwire bundle fabrication design is shown in Fig 2a, enabling different wire core and insulating materials, wire-to-wire spacing and tip shape to be created (Fig S3e,f). The central concept is to bundle together insulated microwires using a sacrificial coating on each one to define the wire-to-wire separation. Insulated microwires were prepared either by depositing a glass or polymer coating, or by thermally drawing a glass/metal wire^29,36,37^. We demonstrate bundles made with metal wires of diameters 5–25 µm and wire materials including Au, W, PtIr and PtW. In principle, this strategy should be applicable to virtually any wire material or size. The high conductivity of metals allows for ultrathin metal cores (down to <1 um), with an adjustable insulating layer from <1 µm to >50 µm thick. In typical preparations, the bare metal wire was wrapped around an open 10 cm spindle, followed by depositing a glass insulating layer ∼1 µm thick using silane decomposition in a Thermco low-temperature oxide furnace at 300 °C, providing a robust inorganic insulating layer^38^.

**Figure 2:**
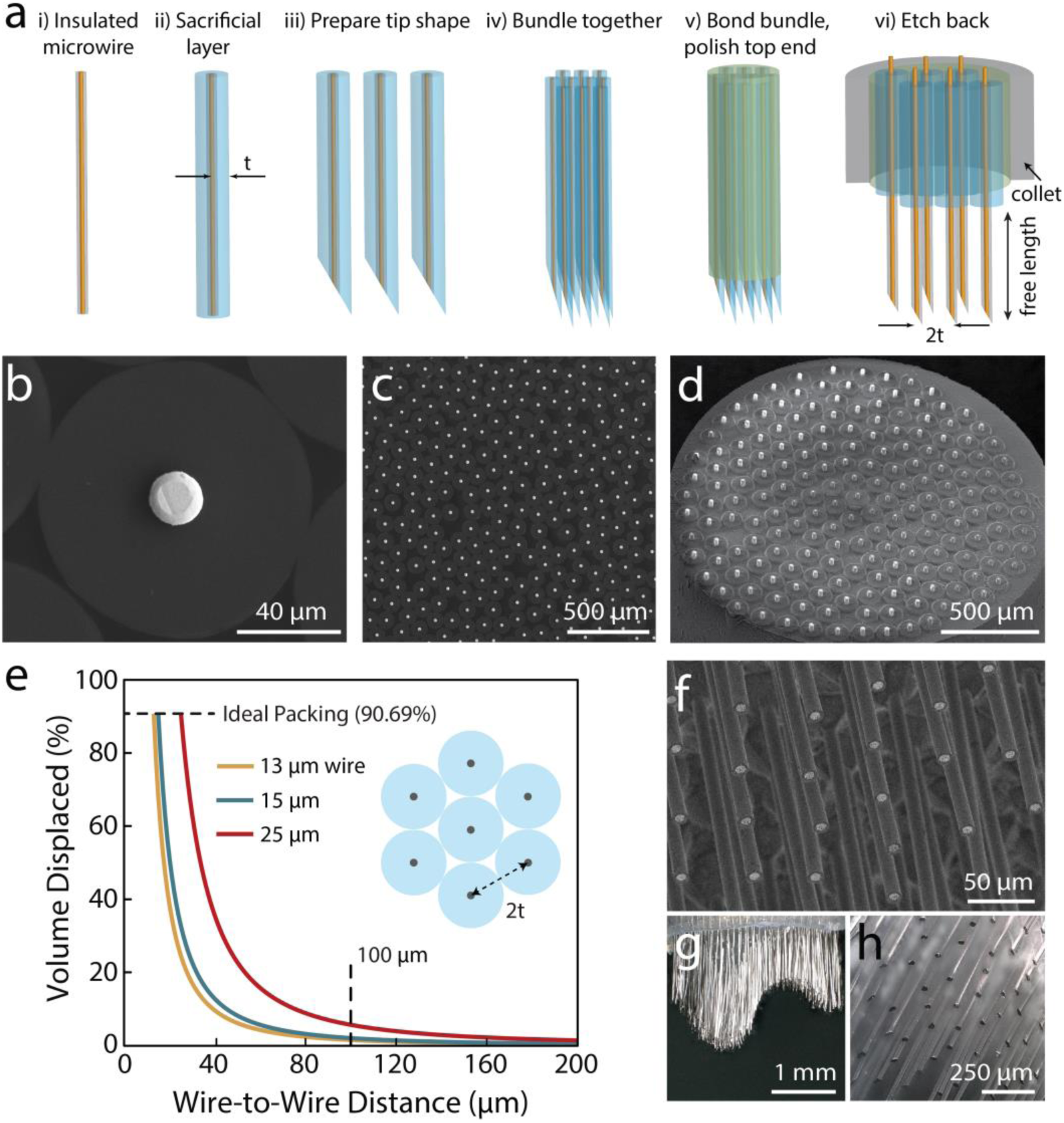
Microwire bundle fabrication. **a**, Fabrication procedure of microwire bundles. (i) Individual microwires are electrically insulated with a robust ceramic or polymeric coating. (ii) A sacrificial layer of parylene-C (PaC) is coated onto the wires to provide spacing. (iii) If desired, the tips of the microwires can be polished to an angular tip, or electrosharpened. (iv) The wires are then bundled together, either by spooling the wire, or mechanical aggregation. The wires naturally pack in a honeycomb array. (v) The bundle is infiltrated with biomedical epoxy to hold the wires together, then the top (proximal) end polished for mating to the CMOS chip. (vi) The proximal end is etched 10-20 µm to mate to the CMOS chip and the distal end of the wires is released by etching with oxygen plasma, allowing each wire to penetrate individually. A threaded collet is added to hold the bundle. **b**, A backscatter SEM of an individual microwire with a PtIr conductive core and 45 µm PaC coating (a, ii). **c**, The wires pack into a honeycomb structure, and epoxy is infiltrated in between to fill the gaps (a, v). **d**, The proximal end of a bundle of 177 20 µm Au wires with 100 µm spacing after etching to expose the conductive wire. **e**, The predicted volumetric displacement of bundles of microwires as a function of wire-to-wire distance, determined by the wire size and sacrificial coating thickness (*t*). This assumes a perfect hexagonal packing fraction of ∼90.69%. For 15 µm wires with 100 µm spacing, the volume displaced is 2%. **f**, The distal end of a bundle of 600 7.5µm W wires coated with 1 µm of glass after etching to remove the PaC and embedded epoxy. **g**,**h**, The distal end (PtW 20 µm wires, 100 µm spacing) can be shaped with single wire precision to simultaneously access different depths in the tissue.

An important aspect of designing microwire bundles for neural recording is controlling the wire-to-wire spacing. Insufficient spacing makes the bundle behave as a solid object during tissue insertion, rather than a collection of individually penetrating microwires. Here, a sacrificial layer of parylene-C (PaC) was deposited via chemical vapor deposition (LabCoater, SCS, IN) onto the glass-insulated wires to set the inter-wire spacing at two times the coating thickness (Fig 2a (ii)). Other materials could be used as well, but PaC was chosen for its chemical inertness, biocompatibility and vapor-phase deposition^39^. Fig 2b,c shows a cross-section from a bundle of 1000 wires 15 µm in diameter with 45 µm thick PaC, giving a center to center wire spacing of 100 µm. This distance could be varied from ∼1 µm to ≥150 µm depending on the desired spacing by altering the thickness of the PaC.

Mechanically bundling the wires was then performed either by a thread spooling tool (Optima 1100, Synthesis, India) commonly used in the textile industry, or by cutting a spool of wire and manually collecting the wires together. The wires were inserted into a biomedical grade shrink-wrap, which compressed the wires together into a honeycomb array, which was infiltrated with a biomedical epoxy (Epoxy Technology Inc.) to create a consolidated structure (Fig 2a (v)). Bundles were then polished and the proximal end was etched to expose 10-20 um of the conductive wire for connection to the chip (Fig 2d). Bundle size could be readily tuned for different applications: from a few hundred wires for mice, where the total size is constrained by the animal model, to >100,000 for high-bandwidth applications in larger mammals. Figure 1b emphasizes the scalability of this technique, showing a bundle of 8640 gold (Au) microwires with 10 µm of glass insulation, spaced at a 40 µm pitch.

After bundling, the microwires were released at the distal end by etching back the PaC coating and biomedical epoxy with oxygen plasma, exposing the wires for tissue insertion. Fig 2e shows the amount of volume displaced as a function of the wire-to-wire spacing. For 15 µm wires spaced at 100 µm, the size and density we used for *in vivo* recordings, assuming perfect packing^40^, the volume displaced is 2%. Fig 2f-h show images of arrays of released wires, displaying uniform diameter and separation. The free length of the wires was controlled by the etching time and was varied from 0.5 mm to well over 1 cm depending on the tissue insertion depth desired. The maximum free length was limited by the material properties and buckling force threshold of the microwires. For example, 18 µm diameter tungsten wires with lengths of > 5 mm are feasible, while for 20 µm gold wires, lengths greater than 3 mm are likely to buckle upon insertion^41^.

Different tip shapes are also possible through polishing or etching the ends of the wires (Fig S3e-g). This was performed by bundling the wires together with a sacrificial polymer and polishing the bundle at an angle (<15–30°) on a lapping wheel. The angle-polished wires were then released and bundled together again with the desired spatial distribution. Alternatively, the wires could be individually electrosharpened^42^ to form a <100 nm radius tip, then bundled together afterwards. Studies measuring the force of microwire insertion into brain tissue have found microwire size and electrosharpening play an important role in the amount of brain dimpling during insertion^41^.

If desired, the distribution of the individual microwire heights can be shaped into three-dimensional structures before the microwires are released from PaC and epoxy. Fig 2g,h show optical images of bundles with variable length wires. These can be produced in arbitrary shapes: for example, to simultaneously access cortical and subcortical regions. The structure is created by machining the epoxy-bonded bundle with a micromill (Minitech-GX MicroMill) or other tool into the desired shape.

### Microwire Bundle to CMOS Interface

A key aspect of the design is developing a high quality microwire-to-CMOS contact. This is a non-trivial problem, as either non-planarities of the two surfaces or slight angular misalignment result in poor connectivity. The system must also be mechanically robust for long-term operation. Here, we overcame these challenges by revealing part of the bare metal microwire at the proximal end of the bundle (Fig 3a,b) then mechanically compressing against the CMOS chip (Fig 3b, Fig S3). Upon compression, the wires bent over (Fig 3b) and ‘crimped’ against the surface of the chip. Upon release, an imprint of the pixel can be seen on the wire core (Fig 3b, top-right inset, Fig S3c). The height of the wires overcame non-planar structures on the CMOS and subtle surface height variations, while the mechanical crimping provided a robust mechanical attachment.

**Figure 3:**
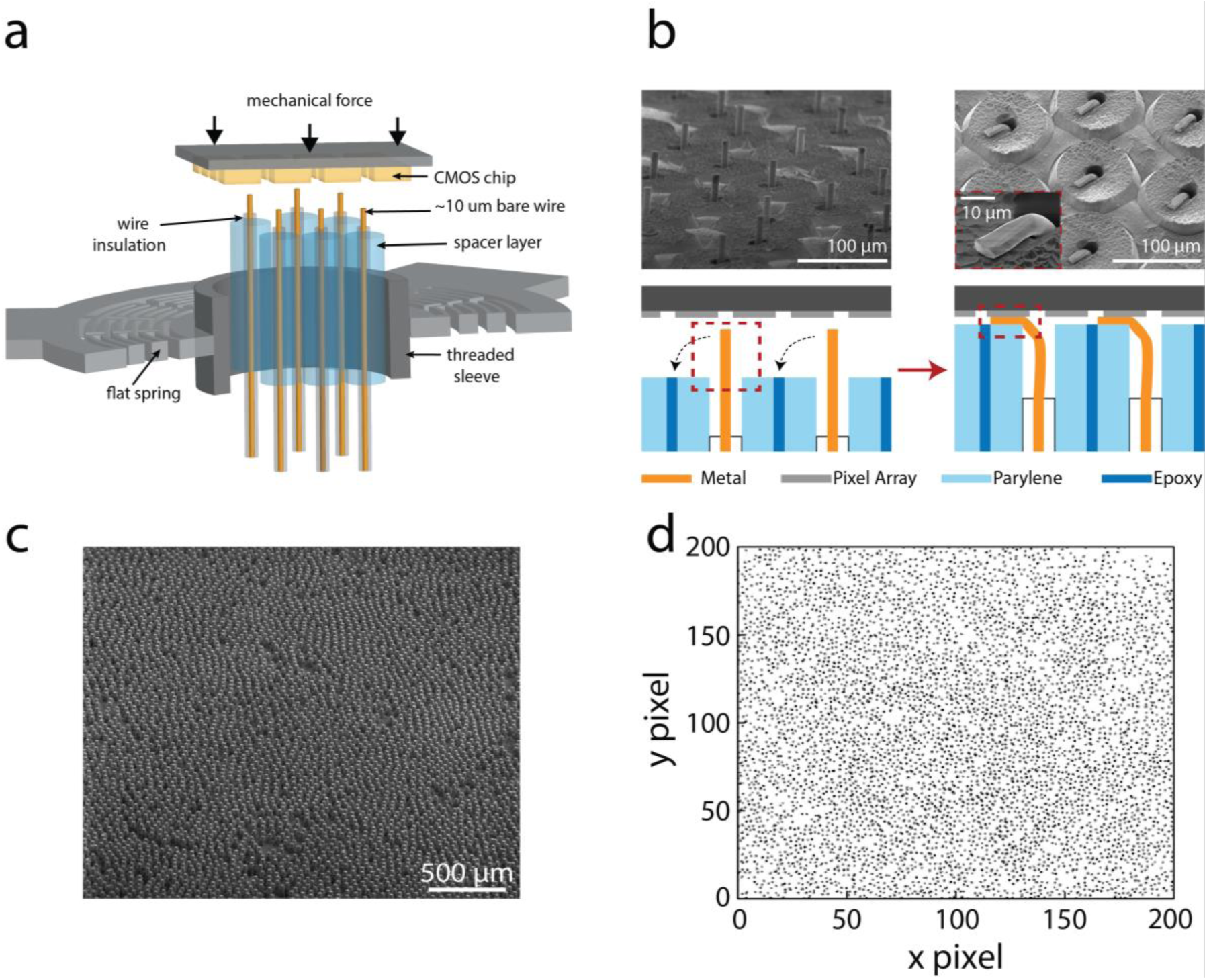
Microwire bonding to CMOS. **a**, Design schematic. The microwires are held by a flat anisotropic spring with a threaded sleeve, and mechanically lowered onto the CMOS chip until high connectivity is achieved. **b**, Prior to bonding, the insulation of the distal end is etched back, leaving ∼10-20 um of bare metal wire. After mechanically pressing to the CMOS chip, these bare wires are mechanically crimped, allowing good electrical contact even with non-planar pixel surfaces. **c**, An SEM focusing on ∼4500 wires of the 8640 bundle shown in Fig 1b, with the corresponding connectivity map, **d**. Each black pixel from the IR camera chip shows a pixel electrically connected to a wire. Over 90% wire-to-pixel connectivity was achieved.

The microwire bundles were prepared for bonding by etching back ∼10-20 µm of bare metal microwire at the proximal end of the bundle (Fig 3b). The free length could be chosen to either be less than the pixel-to-pixel spacing to avoid contacting adjacent pixels, or longer to afford multiple pixel measurements of the same wire. The bundle and CMOS together were mechanically pressed together using a passive, self-aligning mechanical press system designed around a parasitic-error-free, symmetric, flexible diaphragm (Fig 3a, Fig S1)^43^. This structure allowed the bundle to move vertically and tilt during contact, but not rotate or slide, allowing self-aligned contact but preventing scratching or abrasion of the surfaces.

The bundle was then gradually lowered into contact with the CMOS array by turning a threaded screw to extend the bundle (Fig S1a,b,e). Upon compression to the pixel array, each wire crimped, plastically deforming (Fig 3b, Fig S3). This established reliable ohmic contact with the interfacing pixel and was highly uniform over the entire chip surface (Fig 3), with all the wires tilting over in the same orientation (Fig S3d). The degree to which the wires plastically deform depended on the material; large deformations were observed with Au and little observed for W, yet both systems gave equivalent connectivity rates (>95%). After mechanical bonding, epoxy resin could be infiltrated to hold the bundle and chip together (Fig 1d), though the mechanical press alone was sufficient to ensure no change in connectivity over two weeks. Furthermore, the interface was robust to mechanical impacts, such as flicking the bundle with a finger.

This process was highly flexible and agnostic to the identity of the chip; we successfully created bundle-to-chip interfaces with the imaging array chip from a Xenics Cheetah camera (327k pixels, 10×10 µm pixel with 20 um pitch), an OLED display chip from Olightek (1.44M pixels, 4×13 µm pixels), and a multi-electrode array device^44^ (25.6k pixels, 17×17um pixels). The sizes and topologies of each of the pixels in these chips was different, ranging from polished flat Au surfaces to windowed pixels with Al contacts (Fig S3a). If desired, Pt contact pads could be lithographically deposited onto the surface of the pixels to increase the electrically active contact area (Fig S3b).

Post-mating, the microwire-to-pixel connectivity yield was measured by the electrical conductivity and noise characteristics of each pixel. Connectivity between the bundle and chip was high, with >90% of the wires contacting a pixel reproducibly. Figure S2g shows a connectivity map of a bundle of 184 wires, with 177 successfully contacted (96% yield). Highlighting the scalability of this process, Fig 3d shows the connectivity of the central region of the 8,640 wire bundle (Fig 1d, Fig 3c; ∼7 mm diameter, 40 µm pitch, 18 µm diameter microwires) mated to an MEA chip (Xenics Cheetah camera) with >90% of the wires connected to an active pixel (Fig 3c,d).

While the microwires are inherently laid out in a hexagonal fashion, the pixels are in a square or rectangular array, which intrinsically presents an alignment challenge. In practice, there are generally many more pixels available than wires, and the need for 1:1 alignment is not an issue. One possible scenario is a single microwire contacts two or more pixels, which provides additional measurements of the same wire. The other scenario, having more than one microwire contacting a single pixel, should be avoided as separation of the two signals is problematic. This situation can be avoided as long as the distance between wires (*l*_*wire*_) is greater than *l*_*pixel*_√2, where *l*_*pixel*_ is the width of the pixels’ active area. As an example, the anticipated connectivity as a function of pixel pitch was calculated from the average number of contacts over all possible lateral and rotation orientations (Fig 4a). We assumed a metal microwire diameter of 15 µm and pixel pitch of 17 µm with a metallic contact area 50% of the per pixel area (fill factor), similar to the experimental conditions. Fig 4a,b shows that 100% of the wires will always contact one or more pixels, and with microwire spacing >18 µm, no pixel will contact more than one wire. This corresponds well with the actual experimental measurements, which found >90% connectivity yields even with no effort to align pixels to wires. Examples of multiple wires contacting a single pixel only occurred for large pixels and pitches greater than the wire-to-wire pitch. Given that modern camera imagers have pixels ∼1–10 µm on a side and displays on the order of 10–20 µm on a side, alignment of the bundles and CMOS is not necessary given our wire pitches (order of 25–100 µm). One possible issue is the metallic contact pads may be recessed or much smaller than the overall pixel footprint. In this case the pixels can be lithographically modified to enlarge the metallic contact region, such as shown in the inset of Fig 4d.

**Figure 4:**
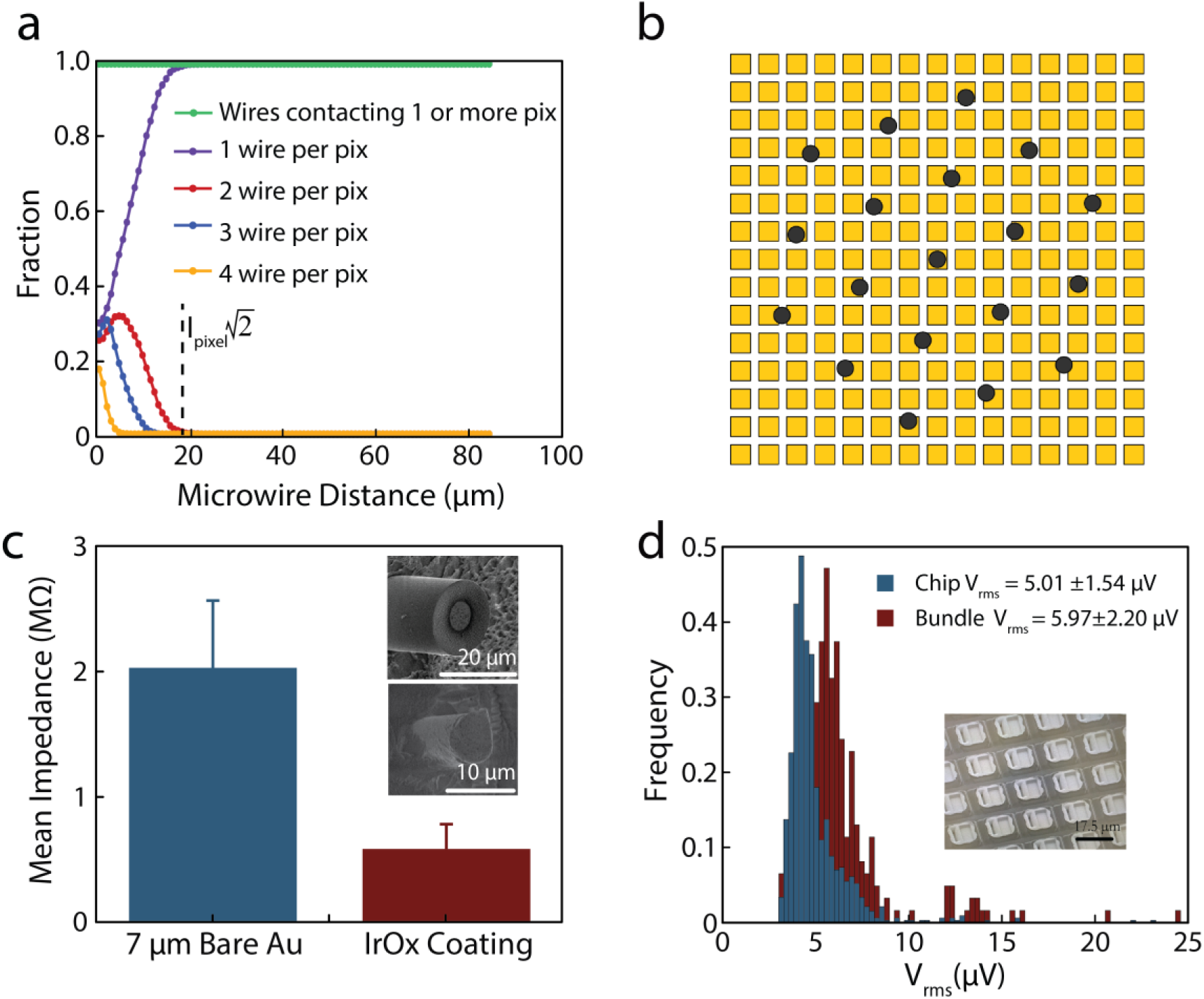
Noise characteristics and wire connectivity. **a**, A hexagonal array of microwires 15 um in diameter can easily achieve 100% connectivity to one or more pixels (green trace) without alignment. Further, no pixels will have more than one wire attached if the wire spacing is greater than l_pixel_√2, where l_pixel_ is the width of the pixels’ active area. **b**, Schematic of the hexagonally arranged microwires overlaid on a square pixel array, showing that alignment is not necessary to achieve unity connectivity yields when the number of pixels is much greater than the number of wires. **c**, Average impedance values of 150 Au microwire bundle before (inset 1) and after in situ electrodeposition (inset 2), measured at 1 kHz. An OLED display chip was used to apply a voltage to individual wires to electrochemically deposit IrOx. **d**, RMS noise of the bare high-resolution CMOS MEA chip (blue), and the same chip after bonding a microwire bundle of 251 PtIr wires, 15 µm in diameter (red). The noise increased from 5.0+/- 1.5 µV (10 Hz – 10 kHz) for the bare chip, to 6.0 +/- 2.2 µV after addition of the bundle, showing the bundle and the mating process did not introduce significant noise. (d, inset) Scanning electron micrograph (SEM) of Pt deposition on the CMOS array to improve electrical contact.

### Electrical performance and *in situ* electrodeposition through CMOS

The electrical characteristics of the microwires were assessed to determine if the mating process increased the recording noise in the CMOS sensor array. First, the impedance properties of individual wires of four different metals (Au, PtW 92/8, PtIr 90/10, and W) were characterized between 0.1–1MHz in PBS. Tests of individual wires found impedances of <1 MΩ at 1 kHz and were strongly dependent on wire material and diameter. This could be substantially reduced by electrodeposition of impedance lowering materials such as IrOx or PEDOT:PSS on the tip of the electrodes^29^. This could either be done by electrically connecting all the wires together and depositing on the array in parallel, or by taking advantage of the wires’ connection to the CMOS to electrochemically deposit on each wire individually. We used a commercial OLED display chip from Olightek to electrochemically deposit IrOx on each wire in a bundle individually, showing a significant reduction in impedance (Fig 4c). The use of the CMOS chip provides a unique method to electrochemically deposit low-impedance materials (IrOx, PEDOT:PSS) on an array of hundreds to thousands of wires simultaneously with control to improve the quality of neural recordings. While all materials tested were capable of recording single-units during acute recordings, PtIr 90/10 was chosen as the preferred material due to its low-impedance (200 ± 27 kΩ) and long term biocompatibility^45^.

One possible concern with bonding an array of metal wires to a CMOS device is greatly increased noise. We tested the electrical pixel noise before and after bundle mating to a multielectrode CMOS chip previously demonstrated for *in vitro* measurements^44^. The chip has 26,400 pixels, 1024 of which can be addressed simultaneously at 20 kHz with stimulation from 32 independent sites, and is commercially available (Maxwell Biosystems Inc.). The recording noise on the unmodified pixels is low, 5. 0 ± 1.5 µV RMS (10 Hz – 10 kHz), enabling robust electrophysiological recordings and stimulation. Pt pads were lithographically added to increase the pixel contact area to 50% of the pixel (Fig 3d, inset), providing robust heterogeneous integration (>95% connectivity) for all bundles. After mechanical pressing, the noise was measured on each channel of a bundle of 251 microwires of 15 µm diameter PtIr cores with 1 µm glass insulation, spaced at 100 µm apart (Fig 4d). The RMS noise was ≤ 5.97 ± 2.2 µV RMS, minimally higher than the unconnected chip itself and well within the tolerable range for *in vivo* recordings.

To assess the temporal stability of the microwire bundle–CMOS interface, the bundle was left pressed for 14 days, and a noise measurement was performed every day (Fig S2i,j). No differences were observed in the connectivity and minor fluctuations in the average noise over this period. Collectively, these measurements show that the addition of the bundle to the CMOS added very little noise to the system, even for relatively long electrodes (7.5 cm), and is mechanically and temporally robust. Figure S2 shows the noise characterization for several different microwire bundle materials and configurations, all demonstrating minimal additional noise relative to the bare chip

### Retinal Recordings

To test the ability of the completed device to record neural activity across a planar surface, we used an *ex vivo* preparation of rat retina, where previous efforts at large scale electrical recordings from retinal ganglion cells (RGCs) provide a performance baseline^46–48^. A dialysis membrane held a small piece of isolated retina against the bundle in a perfusion chamber (Fig 5a), then a 138-wire bundle (mated to the high-resolution CMOS MEA^44^) was lowered into contact with the retina. Activity was recorded in response to both steady, ambient light and to a pulsed visual stimulus, and spikes were sorted using standard procedures (*Mountainsort*^49^, see methods). Recorded spikes presented typical single-unit signatures, i.e. a localized detected action potential at one wire with smaller peaks on adjacent wires (Fig 5a). The highlighted spike waveform in each box is a single unit, and the traces around it are the electrical activity on nearby channels at the times that unit fired. Spiking activity was recorded with high yield (total 152 units; 1.1 units/wire) and high signal-to-noise ratio (>4.5x s.d.) (Fig 5b, filtered voltage traces). Increases in spiking activity were observed during the light stimulation pulse, particularly at its onset and offset (Fig 5c,d). This corresponds to the known existence of ON and OFF type RGCs that respond transiently to light steps, evidence that the spikes were indeed RGC activity. These retinal recordings demonstrate the ability of the system to record single units at high data acquisition rates and high signal to noise.

**Figure 5:**
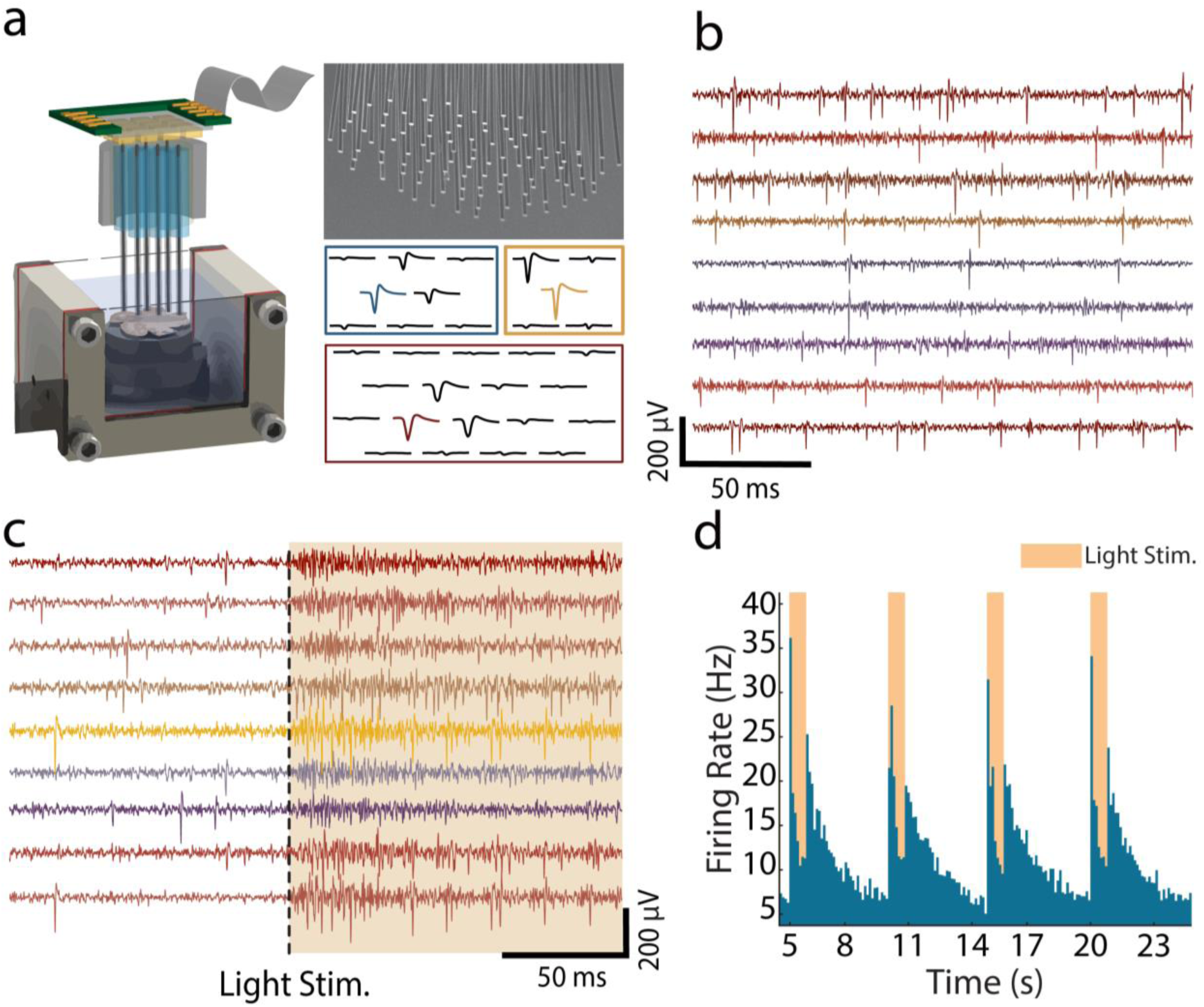
Retinal slice recordings. **a**, Custom-built perfusion chamber used for dissected retina recordings. The retina sits elevated on a dialysis membrane. Sample waveform distribution across a microwire bundle from retinal recordings. **b**, Recordings of spontaneous firing of retinal ganglion cells (RGCs). **c**, Firing of RGCs in response to light stimulation delivered at the time indicated by the dashed line. **d**, A histogram of the average firing rate of all neurons detected, showing an increase immediately following and immediately after light stimulation.

### *In vivo* recording in awake moving mice

Next, we tested whether it was possible to record neural activity in deep cortical and subcortical areas across a large spatial region in rodents *in vivo*. To accommodate the size of the mouse brain, bundles were fabricated to total diameters ranging from 1.75 to 3.5 mm containing 135–251 PtIr microwires (15 µm PtIr core, 1 µm glass coating, length 1–2 mm). The bundles were flat-polished with ∼100 µm spacing between wires to minimally perturb the brain and reduce the likelihood of trauma to the brain and reduced neural activity^50^. Under these conditions, the microwire bundle displaces at most ∼2% of the tissue volume during insertion (Fig 2e). To confirm insertion, we coated the microwires with Dil (1.5 mg/ml) and performed *post hoc* confocal imaging (Fig S4).

Recordings were performed acutely within 2 hours of bundle implantation in deep layers of motor and somatosensory cortices and the dorsal striatum. Mice were allowed to run voluntarily on a spherical treadmill, floated with air pressure, in a head-constrained condition while recording with the bundle and high-resolution CMOS-MEA interface^44^ (Fig 6a)^51^. Spiking activity was readily observed in most of the wires in the bundles across a horizontal layer (Fig 6b). Around a hundred to more than two hundred putative neurons (92–221 neurons, see methods) were reliably identified across a large horizontally extended area in each recording during a typical 5-minute recording session, which yielded 0.56 ± 0.11 (0.3–0.89) single units/wire.

**Figure 6:**
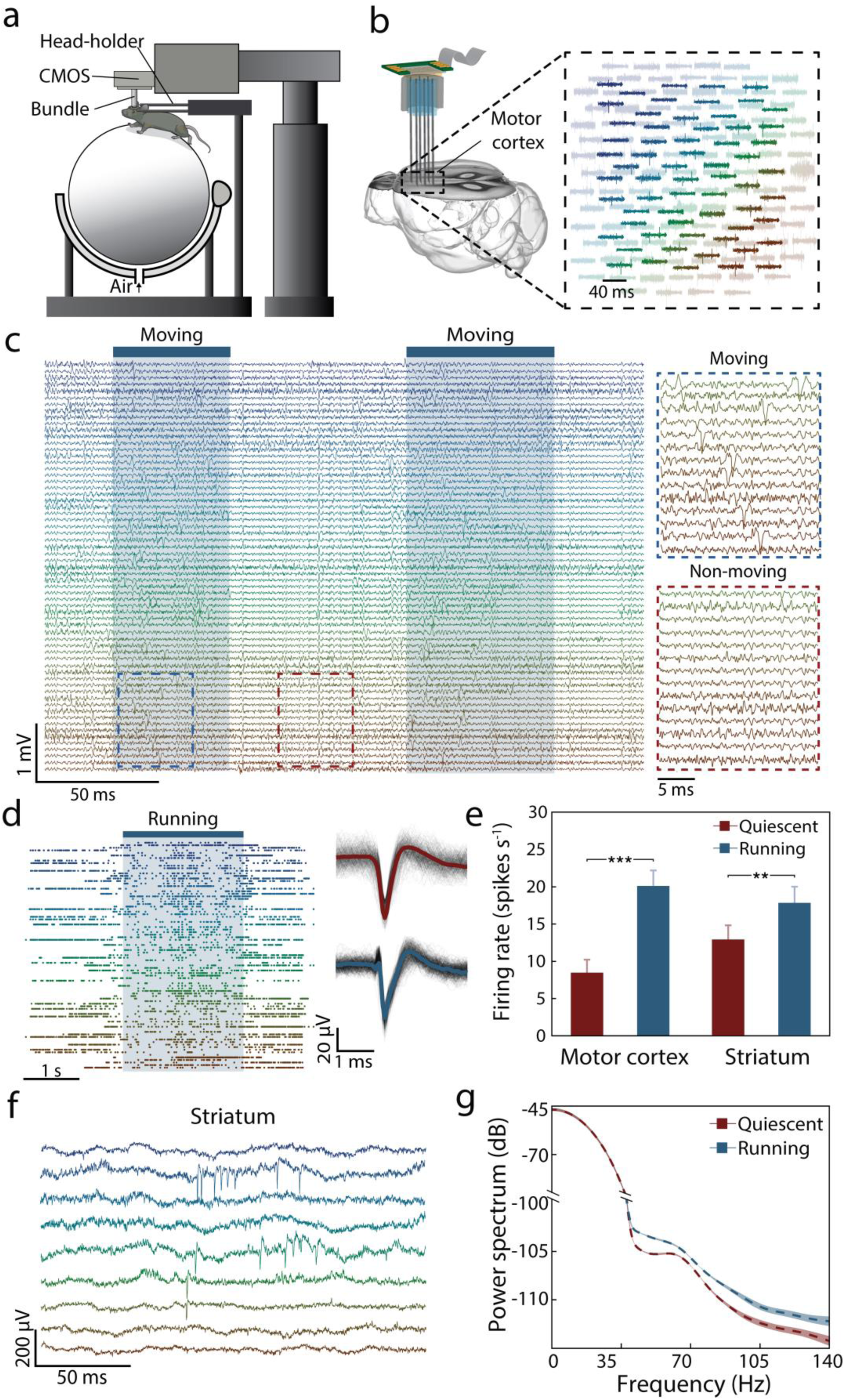
*In vivo* recording in awake moving mice. **a**, A schematic of the *in vivo* recording setup. **b**, *Left*, An illustration of recording across a large spatial extent with a microwire bundle in the motor cortex. *Right*, Representative traces of electrophysiological activity (300 Hz – 6000Hz) from 163 microwires (background traces). Highlighted traces from 67 wires show neural action potentials of a 50 ms snapshot of motor cortical activity during motion. The color code represents the relative positions of the microwires. **c**, Representative traces showing detailed motor cortical activity from the 67 wires highlighted in **b**. The shaded areas indicate the moving episodes of the mouse. Insets show a close look of the representative traces during moving (top) and non-moving (bottom) states. **d**, A raster plot of detected units after spike sorting in a motor cortical recording. Insets show two representative spike-averaged waveforms. Gray traces are 400 randomly selected raw waveforms of two representative detected spikes. **e**, Significantly higher spiking rates were observed during running in both motor cortical and striatal recordings (motor cortex: p < 0.001; striatum: p < 0.01; 30 trials in both areas). **f**, Representative traces (unfiltered) of striatal recording. Both fluctuation in LFP and neural spikes were clearly observed. **g**, The gamma band power was significantly larger during running compared with the quiescent state in the striatum (p < 0.01, 30 trials).

In recordings from motor cortex, spiking activity was highly correlated with motion (Fig 6c,d), consistent with previous findings^52–55^. Significantly higher spiking rates were observed during continuous running sessions compared to the quiescent state (quiescent, 8.4 ± 1.8 spikes/s; running, 20.1 ± 2.1 spikes/s; p < 0.001, n = 30 trials, paired t-test; Fig 6d,e). In a separate experiment, striatal neural activity was recorded with a bundle of microwires with 2 mm of free length in order to access subcortical regions (Fig 6f). Spiking activity of striatal neurons was also correlated with the behavioral states (quiescent, 13.1 ± 1.7 spikes/s; running, 18.1 ± 2.0 spikes/s; p < 0.01, n = 30 trials, paired t-test; Fig 6e). In addition to neural spiking, the rhythmic fluctuations of cortical circuits exhibit strong state-dependent changes^56–58^. We found that low-amplitude LFP fluctuations, especially in the gamma band (30–80 Hz), were significantly larger during free exploration and running states than in the quiescent state (relative gamma band power: quiescent, 0.049 ± 0.012; running, 0.141 ± 0.043; p < 0.01, n = 30 sessions; Fig 6g), consistent with previous reports^59,60^.

Compared to *in vivo* imaging techniques, commonly used for monitoring large populations (hundreds to a thousand) of neural activity, CMOS-bundle devices have several benefits. Conventional two-photon imaging typically records neural activity limited to video frame rates and temporal resolution is further sacrificed when recording from large brain areas^61^. Additionally, imaging techniques are usually limited by tissue scattering and can only be used to record superficial areas without removing brain tissue. For imaging subcortical and deep brain areas such as for GRIN lenses or optical fibers, a portion of the brain tissue is removed^6,61–63^. In comparison, microwire bundles mated to CMOS arrays can record spiking activity from hundreds of neurons and LFPs simultaneously with large spatial extent, albeit with lower spatial resolution, while retaining the benefits of >10 kHz temporal resolution in electrophysiological recordings. In addition, the flexibility of fabricating the length of the distal end of the bundle with single wire precision enables dense recordings from subcortical areas, such as the striatum (Fig 6f), without removing the cortical areas above. Shaping the distal end of the bundle also enables simultaneous access to depths in multiple brain areas. The scalability of this approach is applicable for massive-scale bundle recordings in larger animals, which is particularly challenging for imaging approaches.

## Conclusion

We have demonstrated an effective method to combine the rapid progress in CMOS devices together with brain tissue–compatible probes. Neural recording technologies are rapidly developing, yet are largely based on planar probes (e.g., Michigan Probes, Neuropixels, Neuroseeker), or micromachined silicon arrays (Blackrock Utah arrays). This approach offers a third alternative, retaining the low tissue damage of small microwires, while enabling rapid application of cutting-edge silicon array technology to neuroscience. Additional advances in CMOS technology, such as low-artifact stimulation^64^, higher channel counts^21^ and electrochemical monitoring^65,66^, can be rapidly deployed using this system. Our design is favorable for recording or stimulation experiments that require large area coverage and high density, such as in the visual cortex.

Improvements to minimize the geometrical device and connector form factors are underway, allowing the animal to freely move while recording and/or modulating electrophysiological activity. Subdural implantation may also be possible considering the low power consumption of the neural chip used (< 70 mW)^67^. Due to the control over the depth of each microwire, sampling deep lateral and vertical structures is possible simultaneously with chronic floating microwire-based BMIs, while maintaining ultralow volumetric perturbation of the tissue. The microwire interface provides the link between biological tissue and CMOS electronic technology, enabling the rapid development of silicon-based devices to brain machine interfaces that can readily scale in channel count, temporal resolution, and sensitivity.

## Acknowledgements

Research supported by NIH BRAIN Initiative grant U01NS094248, NIH R21NS104861, NIH SBIR grant 5R43MH110287), DARPA’s NESD program (Contract Number: N66001-17-C-4005), and the Wu Tsai Institute Big Ideas program.

This work was supported by the Francis Crick Institute which receives its core funding from Cancer Research UK (FC001153), the UK Medical Research Council (FC001153), and the Wellcome Trust (FC001153), an HFSP grant to A.T.S. and N.A.M. (RGP 00048/2013), and the Medical Research Council (MC_UP_1202/5). Andreas Schaefer is a Wellcome Trust investigator (110174/Z/15/Z).

The CMOS-MEA chip and setup development was supported by the European Union through the H2020 ERC Advanced Grant “**neuroXscales”** (<u>Contract N° 694829</u>). F. Franke through Swiss National Science Foundation Ambizione Grant PZ00P3_167989

Research at Stanford was supported by NIH BRAIN Initiative grant U01NS094248, NIH R21NS104861, NIH SBIR grant 5R43MH110287, DARPA’s NESD program (Contract Number: N66001-17-C-4005), and the Wu Tsai Institute Big Ideas program. N. Brackbill through NSF GRFP DGE-114747 and NSF IGERT Grant 0801700. This work was also supported by the Francis Crick Institute which receives its core funding from Cancer Research UK (FC001153), the UK Medical Research Council (FC001153), and the Wellcome Trust (FC001153), an HFSP grant to A.T.S. and N.A.M. (RGP 00048/2013), and the Medical Research Council (MC_UP_1202/5). Andreas Schaefer is a Wellcome Trust investigator (110174/Z/15/Z).

The high-resolution CMOS-MEA chip and setup development was supported by the European Union through the H2020 ERC Advanced Grant “**neuroXscales”** (<u>Contract N° 694829</u>). F. Franke through Swiss National Science Foundation Ambizione Grant PZ00P3_167989

## Materials and Methods

### Microwire Bundles

To fabricate insulated microwires in house, PtIr, W, and PtW microwire was purchased of varying diameters (5–125 µm) from Goodfellow Inc. Using a custom spooling rig, microwires were wrapped from their spool onto a custom rack wherein each wire was spaced by ∼500 µm allowing for subsequent chemical vapor deposition (CVD) processing. Silica deposition was done at 300 °C in a low-pressure chemical vapor deposition furnace (Stanford Nanofabrication Facility). A custom rack allowed for >1 km of microwire to be coated at the same time. Using the same rack, PaC was deposited using chemical-vapor deposition (SCS Labcoater 2) to desired thickness, determining the inter-wire separation. Wires are subsequently cut and mechanically bundled together, naturally aggregating into a honeycomb hexagonal array, via shrink wrap. Bundles were then embedded in a biomedical grade epoxy (EpoTek 301, Epoxy Technology Inc.) and cured at 65 °C for 2 hours. Bundles were then placed into borosilicate glass tubes (6 mm outer diameter, 4 mm inner diameter) to aid in handling and sealed using the same epoxy. Polishing of both ends of the bundle was done by successive SiC based grit (Buehler, CarbiMet S, 600, 1000, and 1200), terminating in a hardened silica slurry on a polyester mesh, accomplishing < 10 nm RMS roughness. The bundle ends were then washed with soap, distilled water, and isopropyl alcohol.

Post-polishing, one end of the bundle (proximal end) was dipped in crystalbond (SPI Crystalbond 509) to protect the polished surface. The distal side, the ‘neuronal end’, underwent a piranha etch (3 parts concentrated sulfuric acid and 1 part 30% hydrogen peroxide) for 5 minutes to remove the epoxy embedded between the wires and subsequently washed with soap and distilled water. The bundle was placed in an oxygen plasma to etch away the parylene C on the ‘tissue end’, exposing the glass-ensheathed microwires. The free length of the microwire was determined by the depth of the piranha etch, as the epoxy between the parylene coated wires significantly reduces the etch rate (lateral vs vertical etching). After plasma etching, the wires were cleaned with soap and distilled water. To prepare the proximal end, the crystalbond was removed by placing the bundle in acetone, and placed back in the oxygen plasma briefly to etch ∼1–20 µm of epoxy and parylene between the microwires. The proximal end was then submerged in a 2% hydrofluoric acid etch for 8 minutes to remove the glass coating on the microwires.

To alter tip geometry and/or add a Pt-black coating to lower electrode-electrolyte impedance, microwires coated with glass and PaC were placed into a glass tube, and infiltrated with Apeizon black wax W. Polishing of the wire aggregate occurred at the desired angle, typically done at 24° to produce an acute tip shape. Sputtering of Pt with a high Ar flow rate produced the Pt black tip coatings. Coated microwires were released in toluene, dissolving the binding agent. Subsequent bonding is done as described previously. An alternative process was the use of micromachining the distal end, terminating with the use of a grit-based tooling bit to polish the machined surface (Fig 2g,h). The distal end of this bundle was then etched with oxygen plasma to release the individual wires. The glass layer is unaffected by the oxygen plasma, still providing electrical insulation. The proximal end was processed as described previously.

### Impedance Measurements

The impedance properties of individual wires of four different metals (Au, Pt, PtIr 90/10, W, and Pt-black coated W) were characterized between 0.1–1MHz in 150 mM PBS with a Ag/AgCl reference electrode (Gamry Instruments *Reference 600+)*. Tests of individual wires found impedances of <1 MΩ at 1 kHz, and were strongly dependent on wire material and diameter.

### Passive Mechanical Press System and Microwire Bundle to CMOS Interface

To align two flat surfaces (bundle of microwires and pixel array) such that they are perfectly parallel, ensuring a reliable press, we developed a passive (no active electric components), self-aligning press system. The design is structured around a parasitic-error free symmetric diaphragm flexure seen in Figure S1. The flexible diaphragm (FD) was designed to restrict motion along the X & Y axis, and rotation about the Z axis (yaw; parasitic twist), while allowing for motion about the Z axis, and rotation about the X & Y axis (row and pitch, respectively). This allows two flat surfaces to come into even contact when pressed, while maintaining pressure on the two surfaces as the proportional to the flexible diaphragm’s spring constant. The design of the flexible diaphragm was adopted from Awtar et al., exploiting symmetry to eliminate parasitic twist associated with traditional flexible diaphragm designs when deflected^43^. We show the design of our flexible diaphragm, constructed from spring steel and fabricated via photochemical machining (PCM) (Fig S1c). In this design, 16 peripheral flexure arms are used, despite only 4 being needed to create a symmetrical design. This was done to reduce variation in angular stiffness with in-plane axes passing through the diaphragm center. The hollow center of the diaphragm allows for placement of the microwire bundle (Fig S1b). The spring constant of the FD was defined lithographically by the width of the flexure arms, and thickness of the spring steel used. To minimize material creep, small deflections of the FD were maintained, as it was desirable for some experimental settings to keep the microwire bundle pressed for weeks at a time. The design of the flexible diaphragm also has three emanating appendices, used to grip the FD, and restrict its motion in rotating about the X & Y axis when being pressed into contact. To accurately control the press, a threaded design with a nut was used, allowing the user to slowly screw down the FD and contained microwire bundle until the diaphragm was visibly deflected, or high connectivity was observed. Three tightly specified machined slots were placed onto the male threaded component, allowing for placement of the three emanating appendices of the FD. This prevented rotation of the FD and contained microwire bundle, while still allowing for the screwing mechanism to press the bundle onto the pixel array. To account for the backlash between the male and female threading, an external wave spring was added to constantly keep pressure of the FD & male threaded nut (Fig S1e). In this way, by screwing the nut, vertical displacement of the FD and contained microwire bundle can be controlled. Consequently, the force observed by the chip was dictated by the amount of displacement allotted by the vertical displacement of the FD according to hooke’s law.

The FDs are made via PCM, out of spring steel (0.007” thick). Flexure arm width and spring steel thickness are varied to alter the spring constant of the diaphragm along the perpendicular, z-axis. The mechanical press body was machined out of Al and anodized. The nut was made of brass to prevent material catching. The external wave spring was custom made to fit the geometry of the press system. A collet was machined and glued to the bundle via cyanoacrylate. The FD rested on the end of the collet and was compressed via a corresponding female nut. To bring the microwire bundle into contact with the pixel array, the external nut was rotated, until the flexure arms of the FD were deflected (Fig S1e).

A collet was machined and glued to the bundle via cyanoacrylate. The flexible diaphragm rested on the end of the collet and was compressed via a corresponding female nut. To bring the microwire bundle into contact with the pixel array, the external nut was rotated, until the flexure arms of the flexible diaphragm were deflected. The protruding metallic wires on the proximal end are compressed onto the pixel array using the passive mechanical press system, crimping to establish ohmic contact.

### CMOS Lithographic Modifications

Both the IR ROIC^68^ and the high-resolution CMOS MEA^44^ had recessed contact pads. In the case of the latter, alternating SiO2 and SiN layers were placed to passivate for high-resolution CMOS MEA use. To lithographically pattern the surface of the CMOS, Microposit SPR 220-7 was spun into the surface. Exposure of the desired pixels was done using a maskless aligner (Heidelberg Instruments Inc.). Following development, sputtering of a 5/300/5/30 nm stack Ti/Al/Ti/Pt, followed as Al can be deposited thick without issues of film stress. Liftoff in acetone was done to complete the lithographic modifications. Chips were then wire bonded to the custom PCBs.

### High-resolution CMOS MEA

The high-resolution CMOS MEA was operated as described by previous work^44^. After pressing the microwire bundles onto the CMOS MEA, the distal end of the bundle was submerged in 150 mM phosphate buffer solution (PBS) with a reference electrode of corresponding metallic core material (Pt or W). We applied a 1kHz 1 mV sinusoidal waveform at the reference voltage of the CMOS MEA (1.65V) and scanned through the 26,400 electrodes using a SRS DS360 wave function generator. The electrodes with a wire connected would record the sine waveform, and the pixel position was noted to determine which electrodes to route the channels to (connectivity map). Post establishing the connectivity map, noise was measured by placing the reference electrode (Pt) into the saline bath.

### Infrared Camera Chip

Initial characterization of heterogenous integration was performed with a modified Cheetah 640 CL infrared camera (20 µm pitch, 640 x 512 array; Xenics, Leuven). Upon request, the photosensitive layer was omitted by the manufacturer. In place of standard indium bump pads, lithographic modifications were performed on the top layer of the ROIC (Read Out Integrated Circuit) elevating the electrode pads to optimize for heterogenous integration of microwire electrode bundle to chip array. Sampling frequency is correlated with sampling size, ranging from 1.7 kHz to 200 kHz (largest sampling size to smallest). For 8640 microwire bundle shown in figure 1b, recordings were done at 32 kHz. An SRS DS360 wave function generator was used to apply the input signal. Noise measurements and gain measurements were performed as described in the earlier section.

### Animals

Adult (4-6 month) C57BL/6J mice (JAX# 000664) and wild-type Long-Evans rats were used for this study. All procedures were approved by Stanford University’s Administrative Panel on Laboratory Animal Care.

### *Ex vivo* rat retinal recordings

Eyes were enucleated after decapitation of deeply anesthetized wild-type Long-Evans rats, in accordance with institutional guidelines for the care and use of animals. Immediately after enucleation, the anterior portion of the eye and vitreous were removed in room light, and the eye cup was placed in a bicarbonate-buffered Ames’ solution (Sigma, St. Louis, MO). Under infrared illumination, pieces of retina 3–5 mm in diameter were isolated from the sclera and placed ganglion cell side up on dialysis membrane in the perfusion chamber. The microwire bundle was lowered into the chamber until it made contact with the retina, holding it in place against the membrane. The preparation was perfused with Ames’ solution bubbled with 95% O_2_, 5% CO_2_ and maintained at roughly 30° C and pH 7.4 using an in-line heater. Spontaneous recordings were performed under low, ambient light, and pulsed stimulation consisted of a white, full-field pulse delivered with a hand-held flashlight lasting roughly 1 second, followed by roughly 4 seconds of darkness. The recorded data was filtered (bandpass 300-6000 Hz), and spike sorting was performed using *Mountainsort* as discussed in the data processing methods section.

### Mouse surgery and *in vivo* recordings

Mice were anaesthetized with isoflurane and positioned in a stereotaxic frame. Three skull screws were implanted to provide mechanical stability and for use as ground and reference electrodes. Head-plates made of titanium were centered to the intended recording site on the right hemisphere and fixed to the screws and skull with C&B METABOND Cement (Parkell). A 3 - 5 mm craniotomy was made over the recording sites (motor cortex: 1 mm AP, 2 mm ML, 1.8 mm DV; somatosensory cortex: -1.5 mm AP, 2.5 mm ML, 1 mm DV; dorsal striatum: 1 mm AP, 2 mm ML, 3 mm DV). Dura was carefully removed to facilitate bundle insertion (Figure S4a). The mouse was then transferred to the experimental apparatus and allowed to recover from anesthesia. The head mount was positioned on top of a floating styrofoam ball to allow for movement as described in Bennett *et al*^51^. The bundle was inserted through the craniotomy manually, piercing the pia mater. After slow insertion to the final depth, electrophysiological activity was acquired within 2 hours.

The high-resolution CMOS MEA operates at a floating voltage of 1.65 V. Consequently, the animal was isolated from ground (electrically ‘floating’) by connecting the reference of the chip to the skull screws. The bundles used in Figure 6b-d and Figure 6f consisted of 138 and 251 microwires, respectively, of 15 µm diameter PtIr core with 1 µm SiO_2_ insulative cladding and 100 µm pitch. Motion classification was verified by simultaneously recorded behavior data, in which the running and quiescent periods corresponded well to the active and non-active states of our classification criterion, respectively.

### Data processing and analysis

Data analysis was performed using custom software written in Python 3.6.3 (Anaconda linux-64 v7.2.0 distribution, Anaconda Inc.) and Matlab 2018a (Mathworks). Clustering was done using Mountainsort^49^, with an event detection threshold set to 4.5 SD. Putative single units were identified using noise overlap and isolation thresholds described by J. Chung, J. Magland, et. al 2017 (noise overlap <0.03, isolation >0.95)^49^, and further confirmed by manual curation. The putative single units typically had a negative peak at the recording site, with smaller negative peaks at adjacent sites. Each putative single-unit was recorded on ∼1.4 sites, which is in agreement with the spacing between the electrode sites.

Very short-duration (<0.5 ms), ‘triphasic’, symmetric waveforms, consistent with axonal spikes^69^, could also be identified with amplitudes of ∼20–40 µV, with ∼4.5 times signal to noise ratio. Interestingly, signals with positive polarity occasionally occur at the same time as large negative spikes on nearby electrodes, but the delay between positive peaks with respect to the dominant negative peak on nearby channels varies between recording sites. This behavior is carefully studied by Bakkum, D. J., Hierlemann, A., et. al in acute mouse cerebellar slices and rat cortical cultures, possibly representing dendritic activity, but further work is needed to substantiate whether this is the same phenomena we observe *in vivo*^15,70^.

## Supplementary Figures

**Figure S1:**
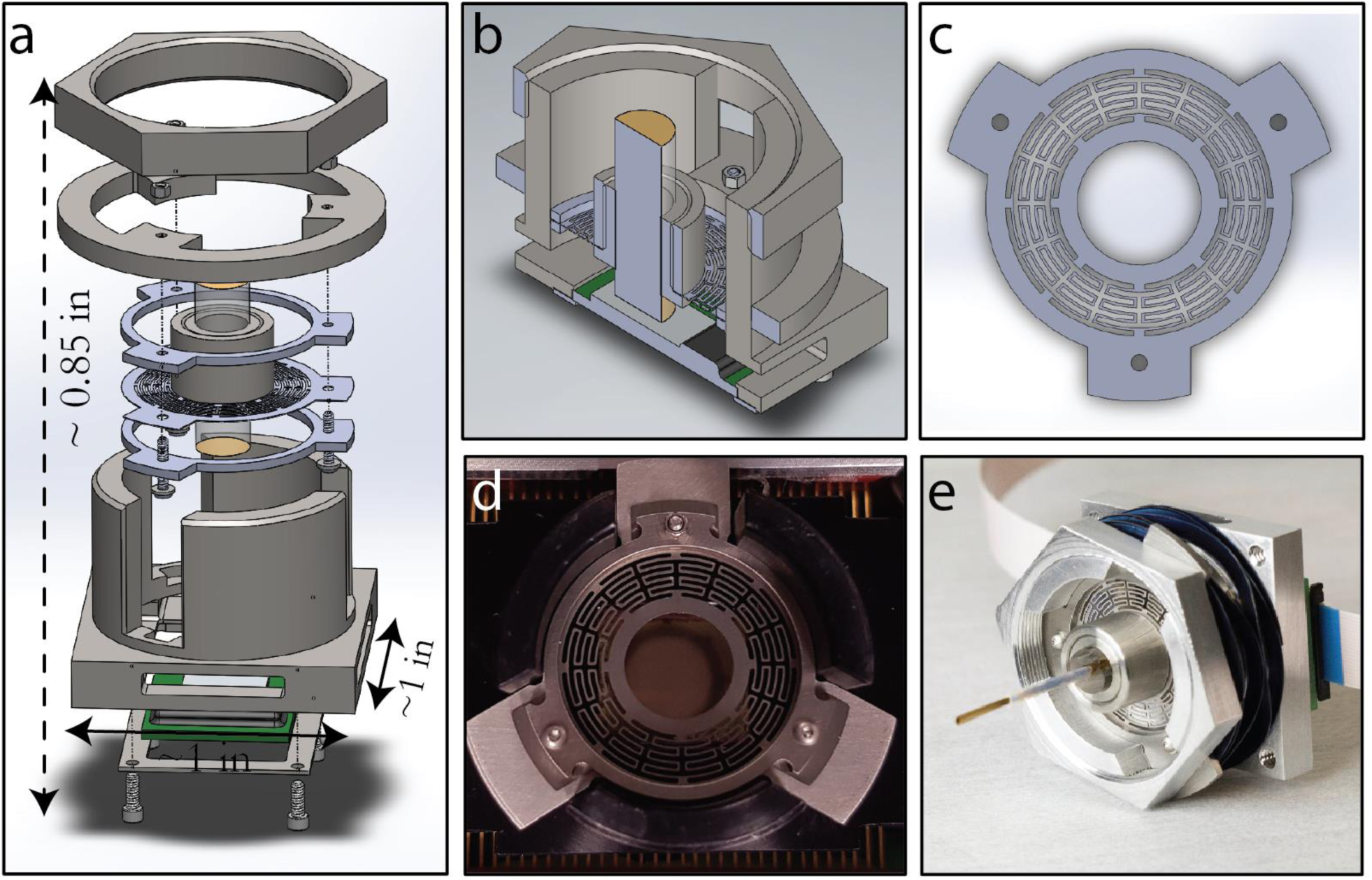
Passive Mechanical Press System. **a**, Exploded image of passive mechanical press system centered with a flexible diaphragm (FD) connected the microwire bundle (portrayed as the glass cylinder). **b**, cross sectional view of press system with the bundle elevated about the CMOS chip. **c**, A CAD drawing of a FD, showing 6 flexure arms and vacant center for bundle placement. **d**, Image of flexible diaphragm over CMOS pixel array (IR ROIC) with bundle removed. **e**, Image of press system post mating of bundle to pixel array.

**Figure S2:**
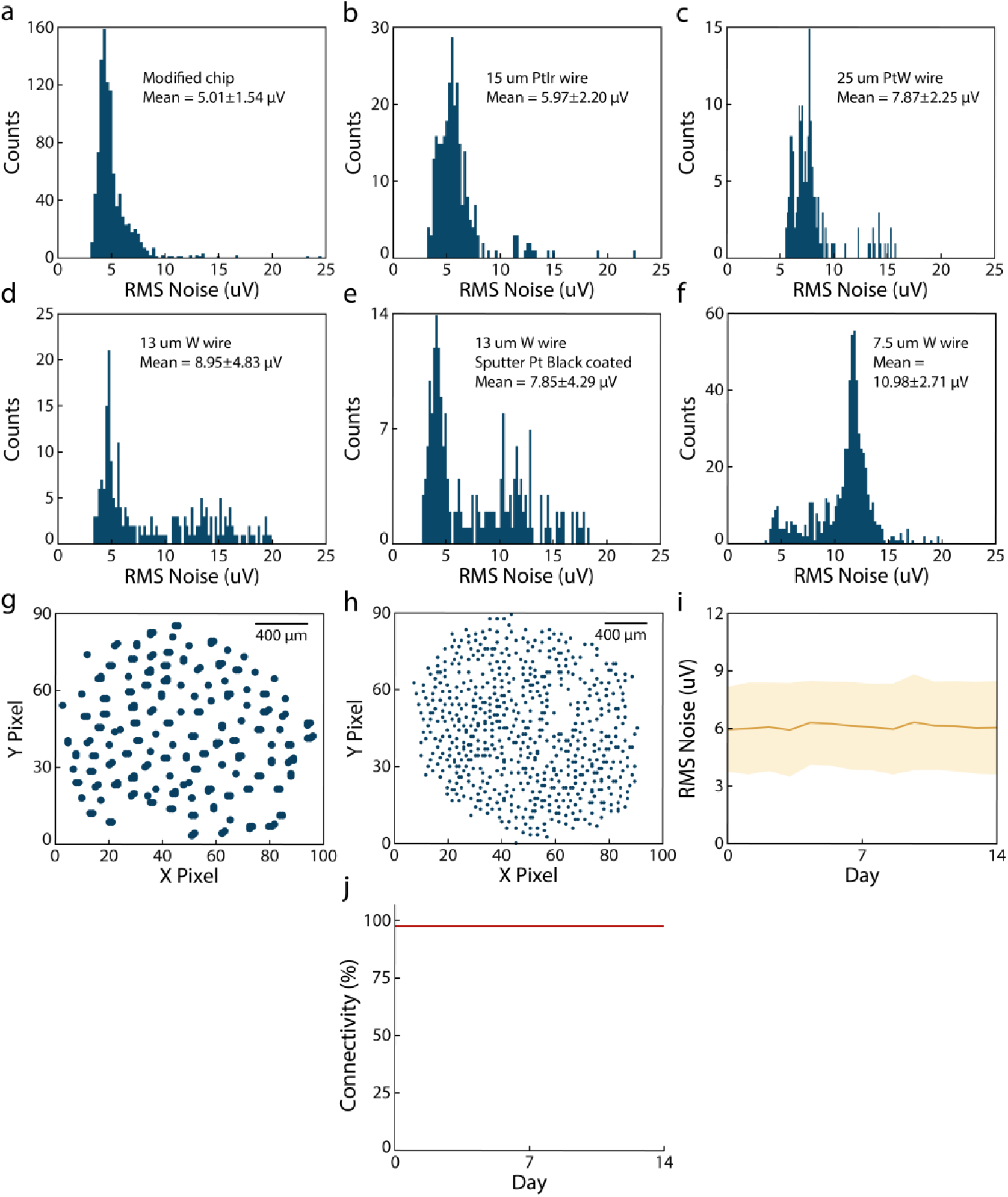
Connectivity maps and RMS noise distribution from bundles made with different materials. **a**, 1024 randomly selected channels from the high-resolution CMOS MEA after lithographic modification without microwires. **b**, 251 15 µm diameter PtIr microwires. **c**, 221 25 µm PtW microwires. **d**, 177 13 µm W microwires and **e**, after Pt-black sputtering on the tips. **f**, 600 7.5 µm W microwires. **g and h**, Example connectivity map from **(d)** and **(f)**, respectively. **i and j**, RMS noise and connectivity of the bundle in **(b)** measured daily for two weeks. The connectivity remained at 97% for the length of the experiment.

**Figure S3:**
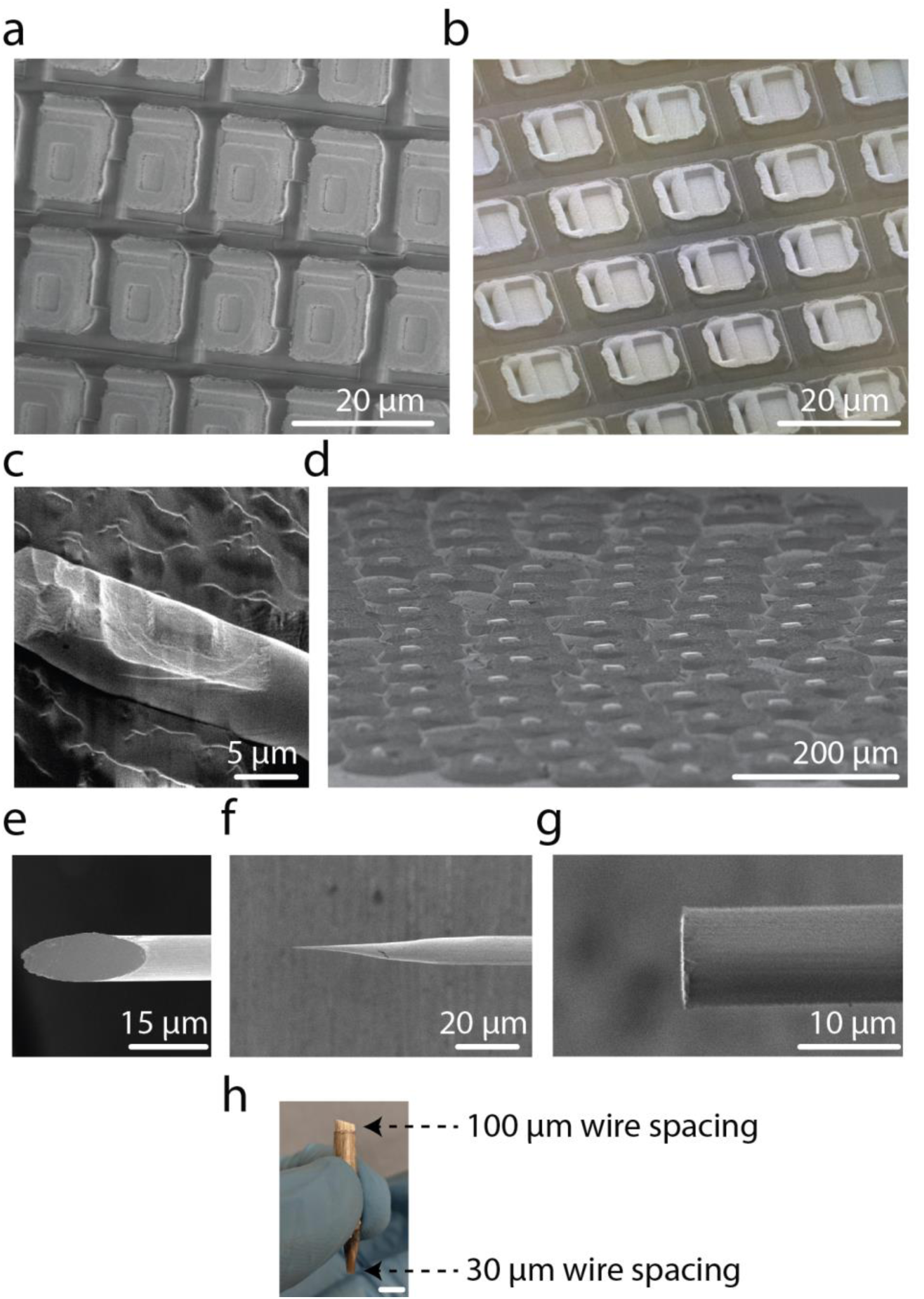
Chip and bundle modifications. **a and b**, SEM image post lithographic modification to elevate recessed contact pads (Cheetah IR chip and high-resolution CMOS MEA, respectively). **c**, An example 7 µm Au wire after pressing onto the chip. The imprint of the pixel **(a)** is seen on the wire. **d**, An example bundle after pressing. All the wires bend in the same direction. **e-g**, Sample possible tip geometries explored (angle-polished, electro-sharpened, flat-polished). **h**, Bundle of PtIr 15 µm wires, wherein the density between the proximal and distal end is modulated independently. The distal and proximal ends have a wire-to-wire spacing of 100 µm and 30 µm, respectively (4 mm scale bar).

**Figure S4:**
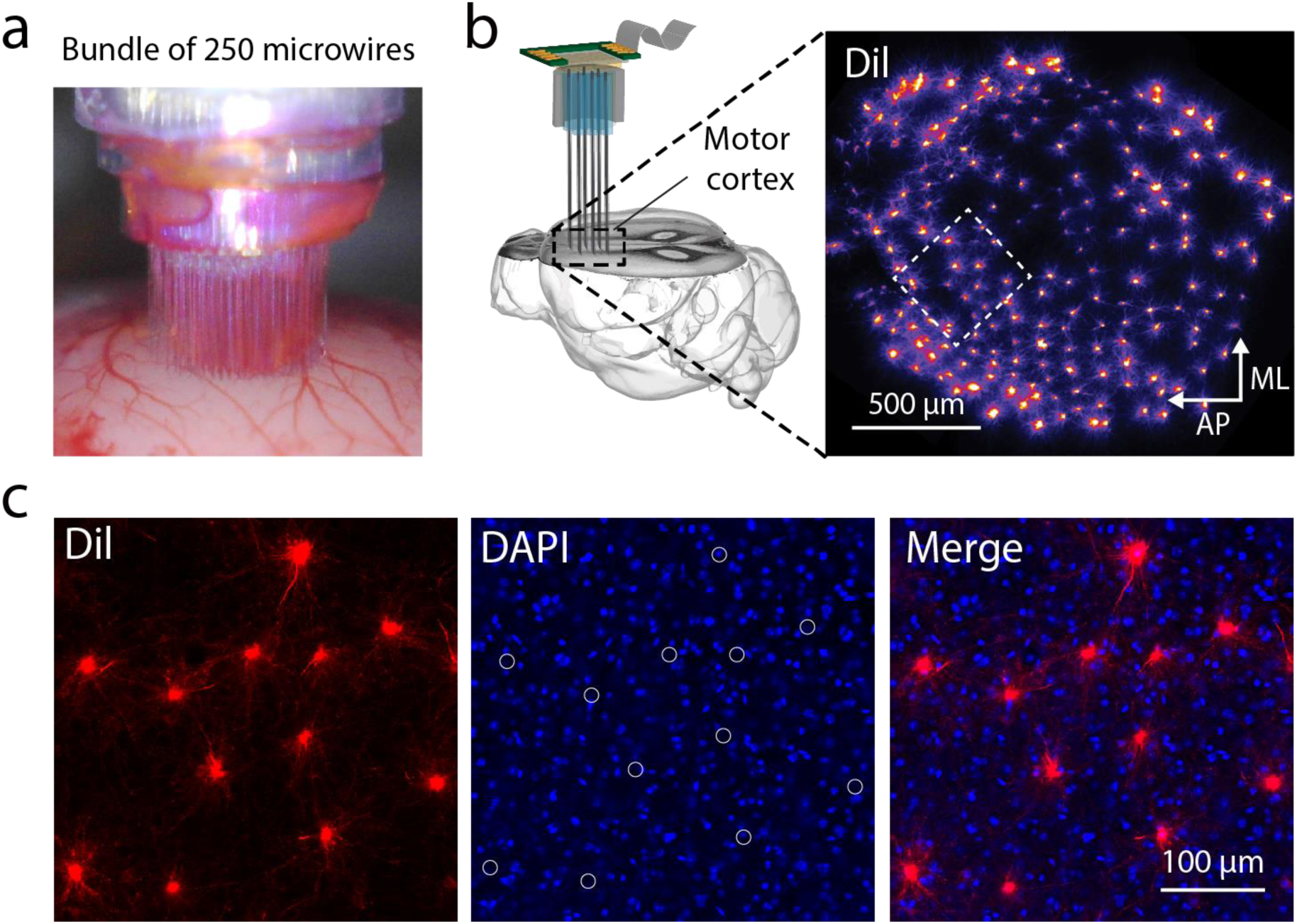
Confirmation of acute insertion of microwire bundle. **a**, A bundle of 250 PtIr microwires (diameter: 15 µm, spacing: 100 µm) coated with Dil (1.5 mg/ml) before insertion into motor cortex of an anesthetized mouse. **b**, A *post hoc* confocal imaging of Dil-labeled cells in a horizontal section at 400 µm depth, confirming the insertion in **a**. AP, anterior-posterior axis. ML, medial-lateral axis. **c**, *Left*, Zoom-in view of the Dil-labeled neurons spaced at around 100 µm, consistent with the space between the microwires of the bundle. *Middle*, DAPI staining of cell nucleus. White circles indicate the location of the Dil-labeled neurons. No clear tissue deformation was observed around the labeled neurons. *Right*, Merge image of Dil and DAPI staining.

